# Impaired acid stress resistance in *Salmonella* Typhi Ty2

**DOI:** 10.64898/2026.04.09.717482

**Authors:** Kiranmai Joshi, Wai Yee Fong, Marie-Pierre Blanc, Fermin E. Guerra, Ferric C. Fang

**Affiliations:** Department of Laboratory Medicine and Pathology, University of Washington, Seattle, Washington, United States of America; Department of Microbiology, University of Washington, Seattle, Washington, United States of America

## Abstract

*Salmonella enterica* encounters acid stress during gastrointestinal transit and within the phagosomal environment of macrophages. Acid stress resistance has been well characterized in *Salmonella enterica* serovar Typhimurium, but comparative studies in the human-adapted *Salmonella enterica* serovar Typhi are limited. We compared the growth of *S.* Typhimurium 14028s and *S.* Typhi Ty2 at pH values ranging from 3-8 and observed that *Salmonella enterica* serovar Typhimurium exhibits enhanced growth at pH 4.5 compared to *S.* Typhi. Comparative transcriptomic profiling of *S.* Typhimurium and *S.* Typhi at pH 4.5 and 7.5 identified numerous differentially expressed acid-induced genes (DEGs), including genes encoding membrane proteins (OmpC, PhoE, HydB), a transcriptional regulator (RpoS), and stress response proteins (YciG, STM14_1829, YmdF). Targeted deletion of selected genes in *S.* Typhimurium significantly suppressed growth at acidic pH, confirming their role in acid stress resistance. These resistance mechanisms are compromised in *S.* Typhi due to pseudogenization. Heterologous expression of pseudogenized genes in *S.* Typhi restored acid tolerance. Collectively, these findings suggest that *S.* Typhi has lost the ability to withstand acid stress due to genomic decay and the loss of multiple genes essential for acid survival in *S.* Typhimurium, reflecting divergent evolutionary paths in these two serovars.

**Importance:** *Salmonella* Typhimurium must adapt to acidic pH conditions in the intestinal tract and the intracellular environment to cause infection. In this study, we show that the enteric fever serovar *Salmonella* Typhi exhibits impaired growth at pH 4.5, in comparison to *Salmonella* Typhimurium. We further show that the loss of specific membrane proteins, a transcriptional regulator, and a family of stress response proteins in *Salmonella* Typhi are responsible for this difference. Collectively, these observations suggest that *Salmonella* Typhi has evolutionarily lost the ability to withstand acid stress due to differences in its interaction with the human host. This has important implications for the pathogenesis of typhoid fever.

## Introduction

*Salmonella enterica* poses a significant global health burden as a leading cause of foodborne illness, resulting in ∼93 million cases of gastroenteritis and ∼14 million cases of enteric fever annually (1, 2). Although most non-typhoidal *Salmonella* infections are self-limiting, severe infections can lead to hospitalization and death, particularly in immunocompromised individuals or those at the extremes of age (3). The ubiquity of this pathogen, its ability to be transmitted between vertebrate hosts and the environment, and the increasing prevalence of antimicrobial-resistant strains have made it a global health priority (4). The disease burden of salmonellosis is highest in low- and middle-income countries, largely due to poor sanitation, but frequent foodborne outbreaks also occur in high-income countries (5).

As a foodborne pathogen, *Salmonella* encounters multiple stresses in the environment and within an infected host, including osmotic shock, high temperature, oxidative stress, nitrosative stress, extreme pH and nutrient deprivation (6). To survive these conditions, *Salmonella* has evolved specific adaptive responses. Resistance to low pH is an adaptive mechanism of enteric pathogens that is crucial for their survival (7, 8). The acid resistance systems AR1, AR2, and AR3 of *E. coli* have been extensively characterized (9). Amino acid decarboxylases play a crucial role in acid resistance of *E. coli* and *Salmonella* by consuming intracellular protons to regulate cytosolic pH (10, 11). Although *Salmonella* is less acid resistant than *E. coli*, it exhibits an acid tolerance response (ATR) involving the regulated synthesis of acid shock proteins, controlled by multiple regulators including RpoS (σ^S^, σ^38^), Fur and the PhoPQ two-component system (12–14). The *Salmonella* ATR can be triggered by modest reductions in pH, which subsequently protects bacteria from more severe degrees of acid stress (15). Although earlier observations suggest that the non-typhoidal serovar *S.* Typhimurium and the enteric fever serovar *S.* Typhi may differ with respect to the ATR and extreme acid stress tolerance (pH 3.3) (16), the mechanistic basis for these differences has not been investigated.

*Salmonella enterica* serovars are categorized as typhoidal or non-typhoidal, a distinction based on host range and clinical characteristics. Most non-typhoidal *Salmonella* serovars (e.g., *S.* Typhimurium, *S.* Enteritidis) infect a broad range of vertebrate hosts and cause localized gastrointestinal disease in humans and systemic infections in animals, whereas enteric fever serovars (e.g., *S.* Typhi, *S.* Paratyphi) are restricted to human hosts, in which they cause systemic infections (17). In contrast to the self-limiting nature of non-typhoidal *Salmonella* gastroenteritis, typhoid fever necessitates antimicrobial therapy, which has been complicated by the emergence of drug-resistant strains (2, 18). Despite sharing over 90% genomic identity, non-typhoidal and typhoidal serovars of *Salmonella* are distinguished by numerous genetic differences, including pseudogenes, unique genes and horizontally-acquired genes (19, 20). Both non-typhoidal and enteric fever serovars are anticipated to encounter acid stress during the host-pathogen interaction: within the stomach, in the distal intestine, and within the phagolysosomes of professional phagocytic cells.

In the present study, the growth of non-typhoidal *S.* Typhimurium 14028s and typhoidal *S.* Typhi Ty2 was compared across a range of pH conditions (pH 3-8). RNA-Seq was performed to analyze the differential gene expression in *S.* Typhimurium 14028s and *S.* Typhi Ty2 at neutral (7.5) or acidic (4.5) pH conditions. Differentially-expressed genes (DEGs) and pseudogenes were analyzed to determine their contribution to the acid resistance phenotypes of *S.* Typhimurium and *S.* Typhi.

## Results

### *S.* Typhi exhibits impaired growth at pH 4.5 compared to *S.* Typhimurium

Growth of *S.* Typhimurium or *S*. Typhi was monitored by measuring the optical density at 600 nm every 15 min in LB broth buffered at pH values ranging from 3-8 at 0.5 intervals over a 12 h period on a Bioscreen C (Growth Curves Ltd, Finland). No growth of either serovar was observed at pH 3.0, 3.5 or 4.0 (Fig. 1A-C). Growth at pH 4.5 was significantly attenuated for *S.* Typhi compared to *S.* Typhimurium, whereas no significant difference in the growth of the two serovars was observed at pH 5-8 except for mild suppression of *S*. Typhi at pH 5.0 (Fig. 1D-K).

**Fig 1.**
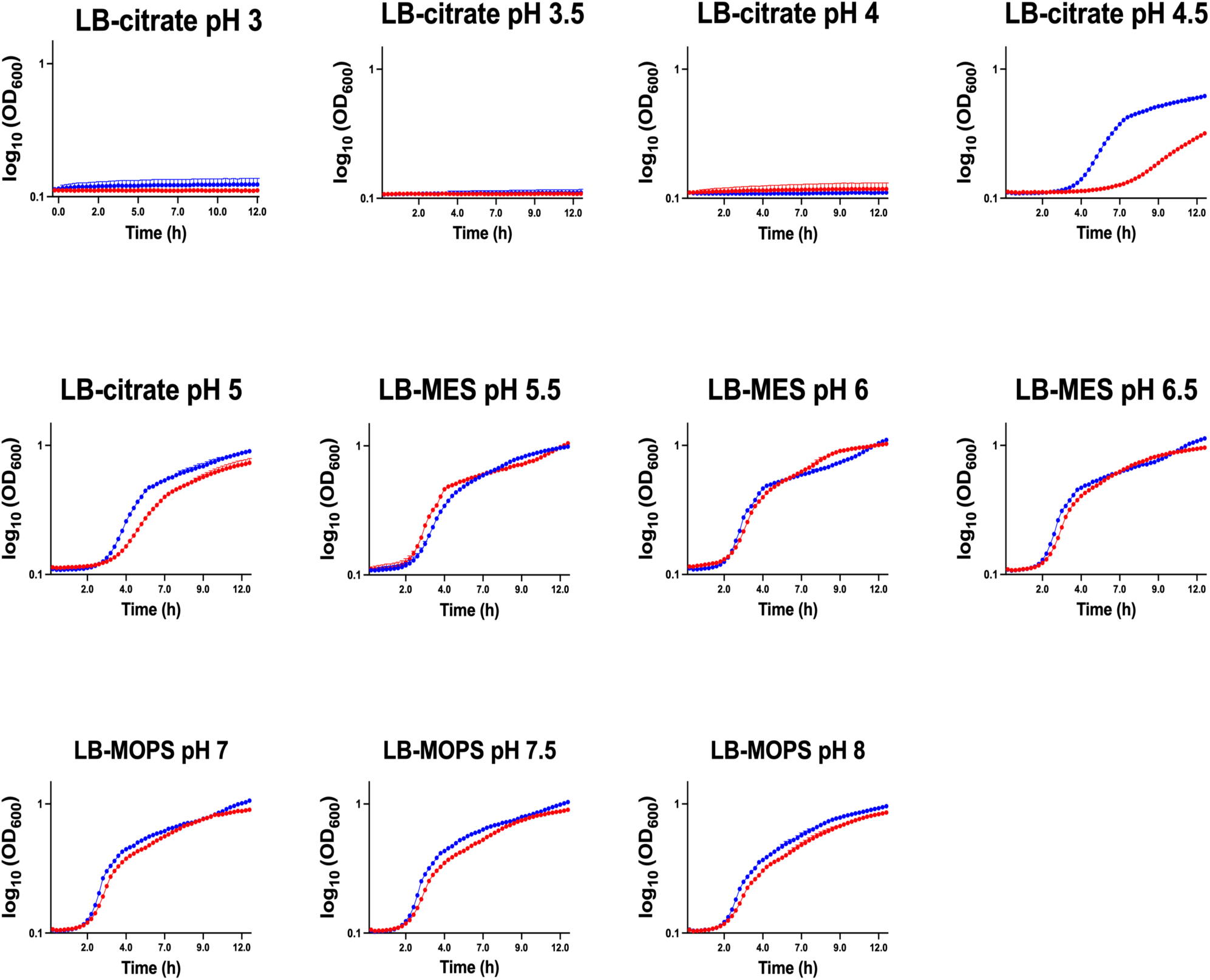
Growth of *S*. Typhimurium and *S*. Typhi in buffered LB media. Optical density was monitored for *S*. Typhimurium 14028s (STm-Blue) and *S*. Typhi Ty2 (STy-Red) grown at various pH conditions. Overnight cultures of STm and STy were diluted to 0.001 OD_600_ in LB buffered with citrate (for pH 3.0, 3.5, 4.0, 4.5, 5.0), MES (for pH 5.5, 6.0, 6.5) or MOPS (for pH 7.0, 7.5, 8.0) and OD_600_ measured every 15 min at 37°C for 12 h in a Bioscreen C.

### Numerous genes are differentially expressed in *S.* Typhimurium and *S.* Typhi at pH 4.5

To identify the genetic differences responsible for greater resistance of *S.* Typhimurium to acid stress at pH 4.5, *Salmonella* gene expression was compared at pH 4.5 and 7.5 by RNA-Seq. Total RNA was isolated from *S.* Typhimurium 14028s or *S.* Typhi Ty2 at early log phase (OD_600_ 0.3) under acidic (4.5) and neutral (7.5) pH conditions, and RNA-Seq was performed. Transcripts were quantified using the FeatureCounts read-counting program. Annotation of transcriptional profiles against the GenBank database yielded 4,849 genes from *S.* Typhimurium, and 4,704 genes from *S.* Typhi. Comparison of gene expression in *S.* Typhimurium 14028s and *S.* Typhi Ty2 at acidic and neutral pH revealed multiple differentially expressed genes at pH 4.5 compared to pH 7.5, including both distinct and common sets for each serovar. A total of 645 genes and 632 genes were upregulated and a total of 665 and 831 genes were downregulated in *S.* Typhimurium and *S.* Typhi, respectively, at pH 4.5 relative to expression at pH 7.5 (log2FC >1, adj-p <0.05). However, considerable differences were observed between the serovars, with 259 *S.* Typhimurium genes upregulated at pH 4.5 relative to *S.* Typhi. The volcano plots and the Venn diagrams in Figure 2A-D represent the number of differentially expressed genes in *S.* Typhimurium and *S.* Typhi at acidic pH 4.5.

**Fig 2.**
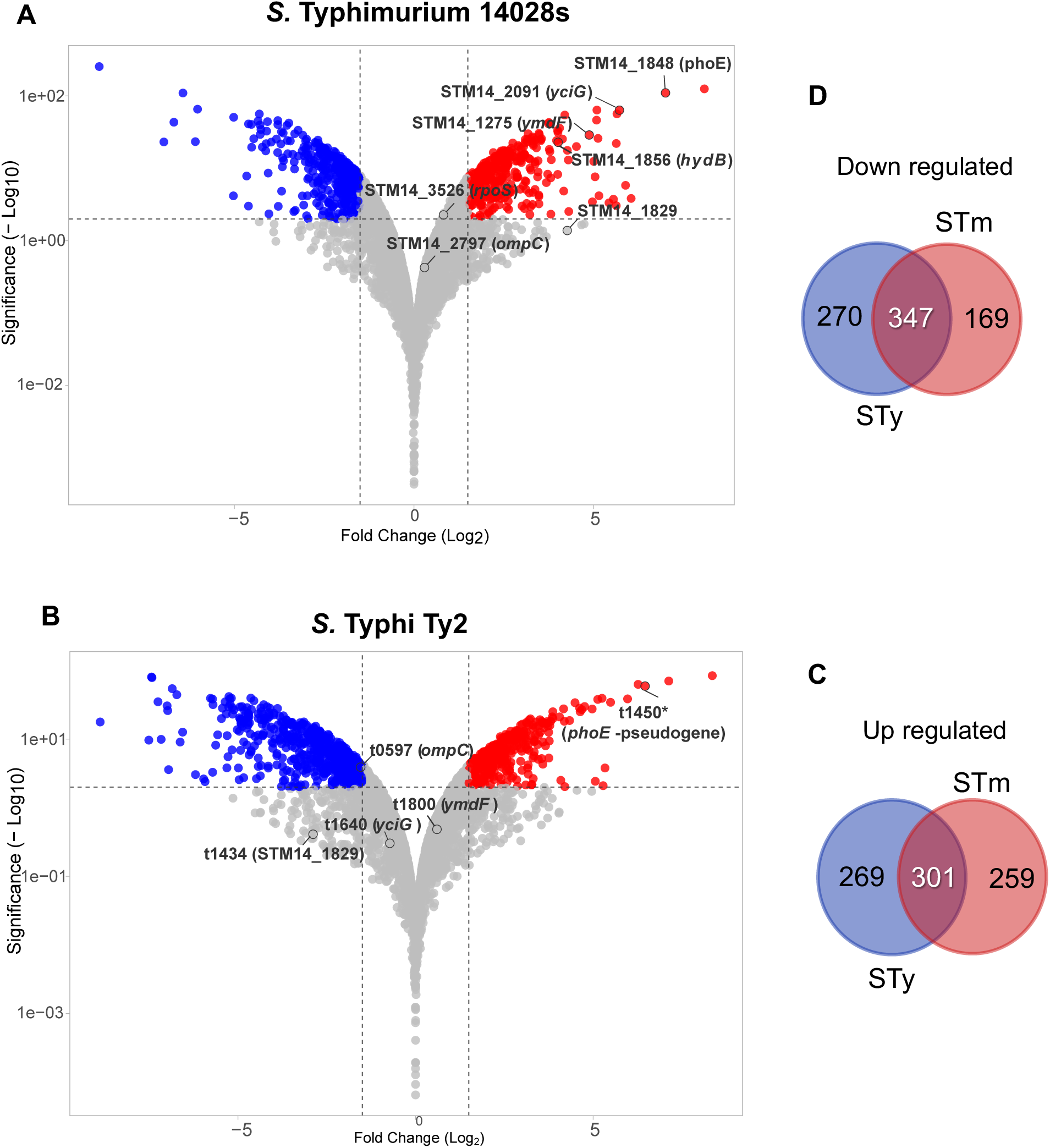
Transcriptomic analysis of *S.* Typhimurium and *S*. Typhi. Gene expression was analyzed by RNA-Seq in *S.* Typhimurium 14028s (STm) and *S.* Typhi Ty2 (STy) grown to early log phase (OD_600_ 0.3) in LB medium at pH 4.5 or 7.5. Differential gene expression in both strains was analyzed using the FeatureCounts read counting program. Volcano plots show differential gene expression in STm (A) and STy (B), highlighting the target genes in the current study. Genes significantly upregulated or downregulated at pH 4.5 relative to pH 7.5 (log2FC >1, adj-p <0.05) are indicated in red or blue, respectively. (C&D) Venn diagrams represent the number of upregulated (C) and downregulated (D) genes in STm (red) and STy (blue), respectively.

Some of the genes that were differentially expressed encoded membrane proteins and stress response proteins. Differentially expressed outer membrane protein (OMP) genes included *ompC, ompF, phoE* and *ompD*. Under acid pH conditions, *S.* Typhimurium exhibited upregulation of the general stress response genes *yciG, STM14_1829,* and *ymdF,* which encode small intrinsically disordered proteins (21), whereas these genes were downregulated in *S.* Typhi (Table S1). Expression of genes encoding transcriptional regulators involved in acid stress responses such as *phoPQ, fur, envZ* and *ompR* revealed no difference in expression between the two serovars, with the exception of *rpoS,* which was slightly upregulated in *S.* Typhimurium but is non-functional in *S.* Typhi due to a frameshift mutation (22).

Specific attention was paid to genes encoding enzymes with known roles in acid resistance, including *narWV, fdhN, hydB, adiA, adiC, speC, speF and cadA* (23, 24). A significant upregulation of *narWV, fdhN, hydB* and *speC* was observed in *S.* Typhimurium, whereas these genes are pseudogenes in *S.*Typhi (25). The *adiA, adiC, speF* and *cadA* genes did not show differential expression at pH 4.5 in *S.* Typhimurium and *S*. Typhi (Table S1).

### *S.* Typhi lacks multiple membrane proteins that contribute to *S.* Typhimurium acid stress resistance

Individual deletion mutations were constructed in differentially-expressed genes encoding outer membrane proteins (OMPs) in *S.* Typhimurium. Growth of mutant strains was compared to that of wildtype *S.* Typhimurium and *S.* Typhi at pH 4.5 and 7.5. Mutants containing single OMP gene deletions (STm Δ*ompC*, Δ*ompD*, Δ*ompF* or Δ*phoE*) exhibited growth phenotypes similar to that of wildtype *S.* Typhimurium (Fig. S1A). Double gene deletions (STm Δ*ompC* Δ*ompF*, Δ*ompC*Δ*ompD*, Δ*ompD*Δ*phoE* or Δ*ompF* Δ*phoE*) did not impair the growth of *S.* Typhimurium at pH 4.5, with the exception of the Δ*ompC* Δ*phoE* mutant, which had a detectable growth defect. Mutants containing triple gene deletions (STm Δ*ompC*Δ*ompD*Δ*phoE* and STm Δ*ompC* Δ*ompF* Δ*phoE*) also exhibited slower growth at pH 4.5 compared to the wildtype *S.* Typhimurium (Fig. 3A, Fig. S1B, C). No growth difference was observed between STm Δ*ompC* Δ*phoE* and wildtype S. Typhimurium at pH 7.5 (Fig. S1D). Collectively, these observations indicate that OmpC and PhoE make redundant contributions to acid stress resistance in *S.* Typhimurium. A notable correlation was observed between the growth kinetics and expression of *ompC* and *phoE* in the RNA-Seq data. Expression of *ompC* at pH 4.5 was higher in *S.* Typhimurium than in *S.* Typhi, and *phoE* was not expressed in *S.* Typhi because of truncation of its coding region (Fig. 3B, C). We performed an allelic exchange to replace the non-functional *phoE* gene (*t1450*) of *S*. Typhi with a functional *phoE* gene from *S.* Typhimurium. Growth curves were then analyzed under neutral and acidic pH conditions. The repaired STy Δ*t1450*::*phoE*_STm_ allele partially restored *S.* Typhi growth at pH 4.5, confirming the contribution of PhoE to *Salmonella* acid resistance (Fig. 3D). There was no difference in the growth phenotype of wildtype *S.* Typhi and the STy t1450::*phoE*_STm_ mutant at pH 7.5 (Fig. S1D).

**Fig 3.**
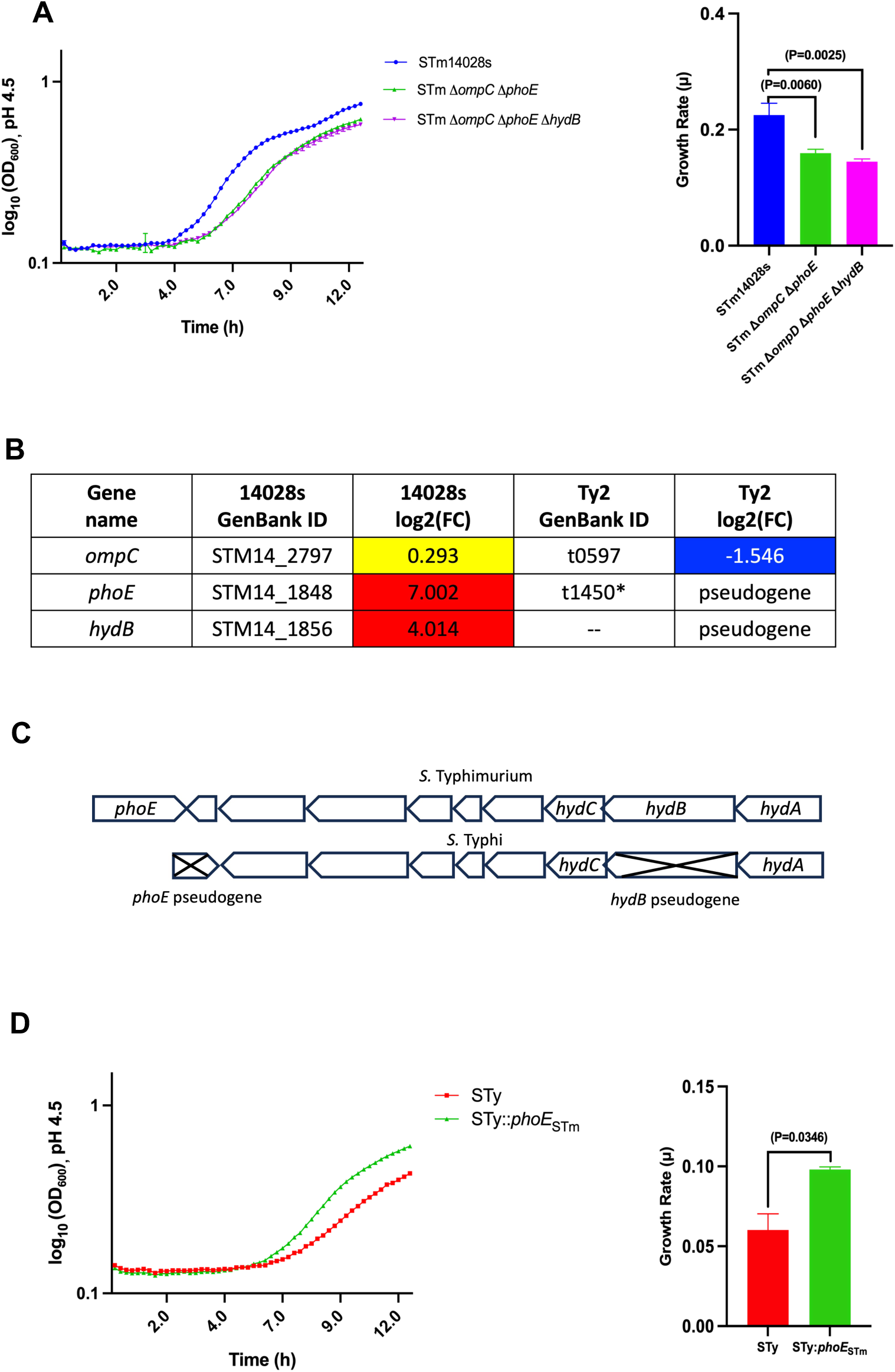
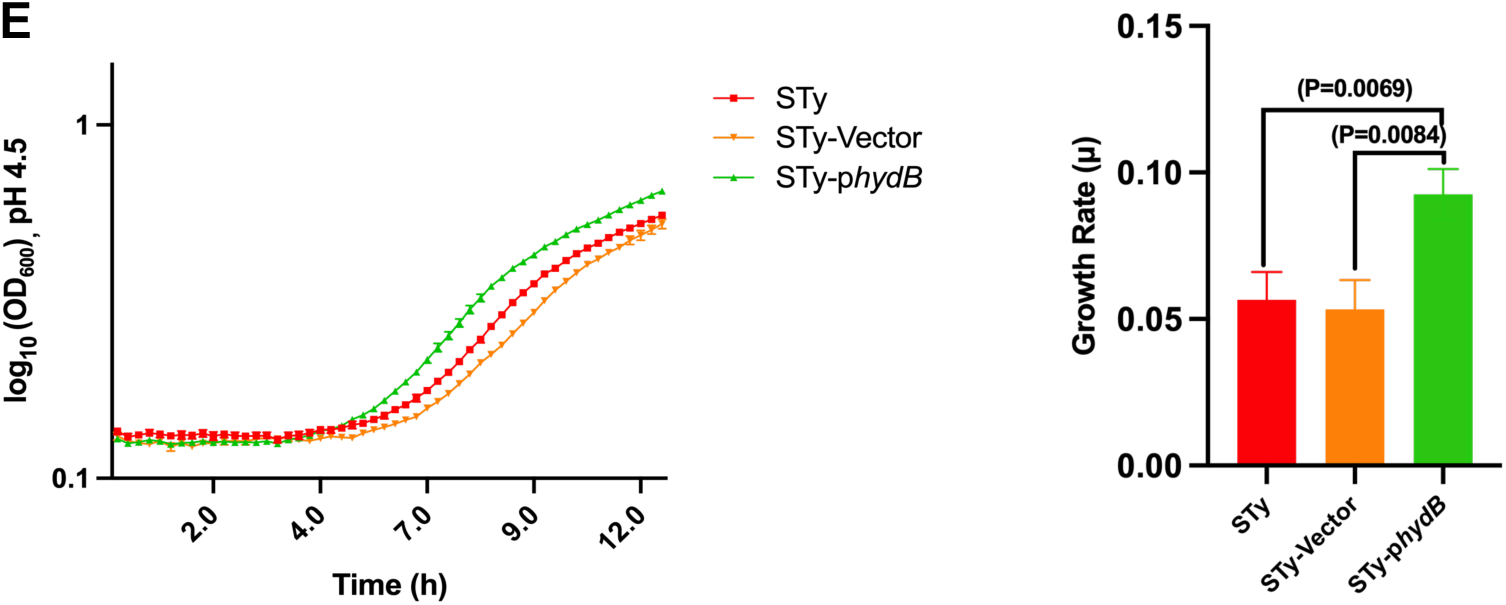
*S.* Typhi lacks multiple membrane proteins that contribute to *S.* Typhimurium acid stress resistance. The porin PhoE and membrane-bound hydrogenase HydB are present in *S*. Typhimurium (STm) but not *S*. Typhi (STy). (A) Growth of wt STm, STm Δ*ompC* Δ*phoE* and STm Δ*ompC* Δ*phoE* Δ*hydB* in buffered LB at pH 4.5. (B) Fold change of OMPs (*ompC, phoE*) and hydrogenase (*hydB*) gene expression in LB buffer at pH 4.5 in STm and STy. (C) Gene maps of STm and STy derived from Biocyc database showing the pseudogenes *phoE* and *hydB*. (D) *phoE* of STm was allelic-exchanged in STy and growth monitored for STm wt and STy::*phoE*_STm_ in buffered LB at pH 4.5. (E) Growth of STm wt, STy-Vector (pJK770) and STy-p*hydB* (pJK770-*hydB*) in buffered LB at pH 4.5. To generate the growth curves, bacteria were cultured in LB-citrate (pH-4.5) or LB-MOPS (pH-7.5) and OD_600_ measured every 15 min at 37°C for 12 h on a Bioscreen C. Typical representative growth curve of three independent experiments are shown. Statistical analysis was performed in GraphPad prism 10.0. Bar graphs represent the mean slopes of exponential growth with error bars representing the standard deviation from three independent assays. *P* values were determined using an un-paired two-tailed *t-*test (A, D & E).

HydB is a subunit of a membrane-bound Hyd enzyme complex that enables *S.* Typhimurium to use molecular hydrogen as an energy source during aerobic respiration (26). Upregulation of *hydB* was observed at pH 4.5 in *S.* Typhimurium, but *hydB* is predicted to be nonfunctional in *S.* Typhi due to a nonsense single nucleotide polymorphism (SNP) (Fig. 3B, C). Although deletion of *hydB* alone had a minimal effect on growth of *S.* Typhimurium at pH 4.5, an additive effect on growth was observed when a Δ*hydB* mutation was introduced into a *S.* Typhimurium Δ*ompC* Δ*phoE* double mutant strain (Fig. 3A, Fig. S1E). Growth of the triple Δ*ompC* Δ*ophoE* Δ*hydB* mutant was unaffected at pH 7.5 (Fig. S1F). Furthermore, heterologous constitutive expression of the *hydB* gene of *S.* Typhimurium in *S.* Typhi on an expression vector enhanced the growth of *S.* Typhi at pH 4.5 compared to vector alone or wildtype *S*. Typhi (Fig. 3E). No difference in growth of *S.* Typhi carrying P*hydB*, *S.* Typhi with the vector alone, or wildtype *S.* Typhi was observed at pH 7.5 (Fig. S1G). Taken together, these results indicate that the outer membrane proteins OmpC and PhoE and the aerobic membrane-bound Hyd hydrogenase contribute to acid stress resistance in *S*. Typhimurium but are absent or deficient in *S*. Typhi.

### *S.* Typhi lacks stress response genes that contribute to *S.* Typhimurium acid stress resistance

A complex network of transcriptional regulators are activated by external stresses such as osmotic shock, oxidative stress, nitrosative stress, nutrient deprivation, and acid stress conditions, which allows *Salmonella* to withstand varying environmental conditions during infection (7). RNA-Seq analysis of *S.* Typhimurium and *S.* Typhi did not show differences in most of the key transcriptional regulators implicated in acid stress resistance, such as *phoPQ*, *fur*, *ompR* or *envZ* (Table S1). However, the *rpoS* gene encoding the alternative sigma factor σ^38^ (RpoS, σ^S^) is non-functional in the *S.* Typhi Ty2 strain due to a frameshift mutation (14, 22) (Fig. 4A). We found that deletion of *rpoS* in *S.* Typhimurium resulted in a severe growth defect at pH 4.5 but not at pH 7.5, consistent with an important role in acid resistance (Fig. 4B, Fig. S2A). As *rpoS* is non-functional in *S.* Typhi Ty2, we investigated the ability of *S*. Typhi CT18, which contains an intact *rpoS* gene, at pH 4.5 and 7.5. Growth of *S*. Typhi CT18 remained defective at pH 4.5, indicating that additional genes deficient in *S.* Typhi are required for acid stress resistance (Fig. S2B, C). In addition, the native *rpoS* gene of *S.* Typhimurium was replaced with the *rpoS* gene from *S.* Typhi Ty2 via allelic exchange. An altered growth phenotype was observed in Δ*rpoS* mutant *S.* Typhimurium and in the allelic-exchanged STm::*rpoS*_STy_ strain at acidic pH, with more rapid entry into exponential growth and reduced terminal OD600, confirming the impaired activity of σ^38^ in *S.* Typhi Ty2 (Fig. 4B and 4C, Fig. S2D).

**Fig 4.**
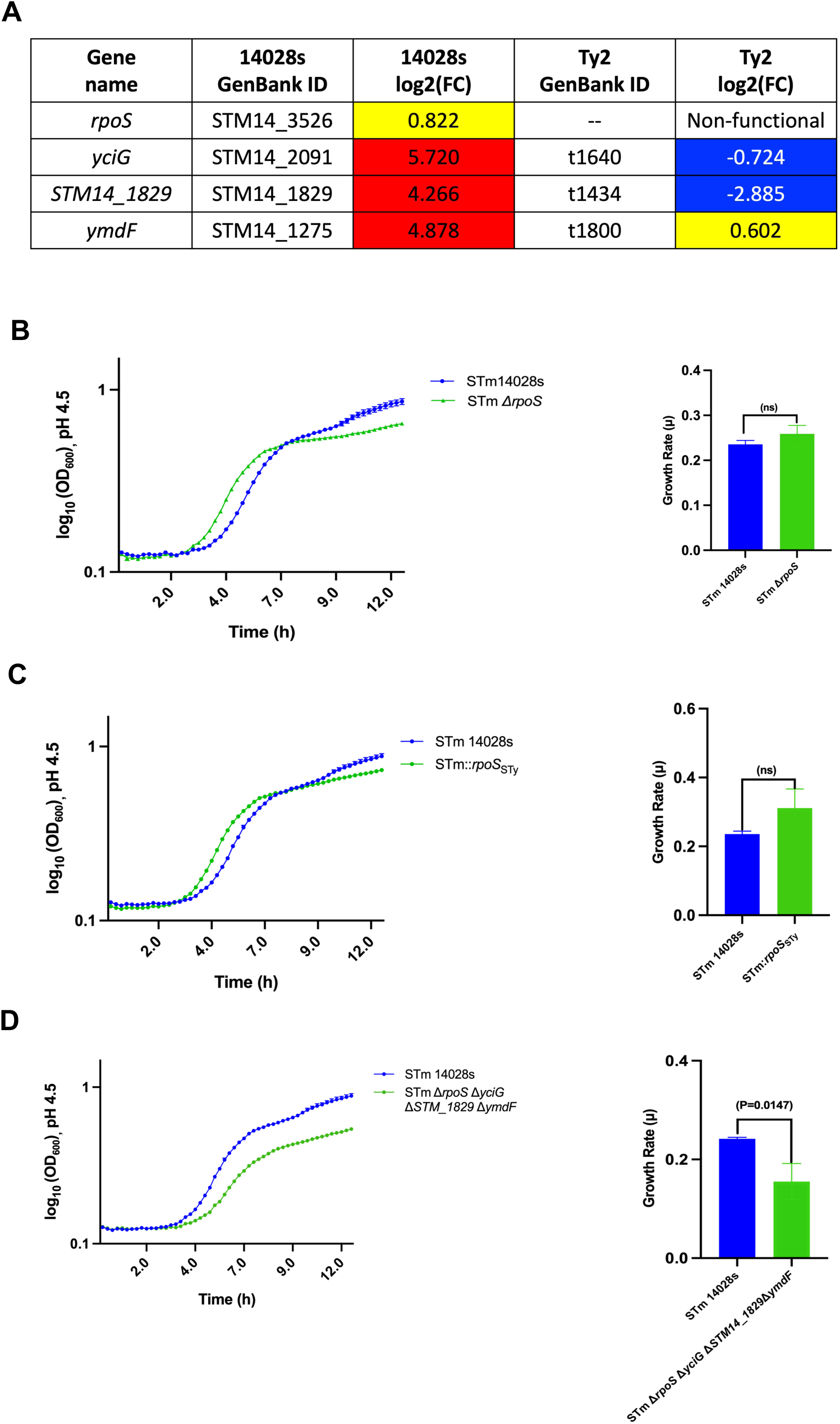
*S.* Typhi lacks stress response proteins that contribute to acid resistance in *S.* Typhimurium. Actvity of the alternative sigma factor RpoS (σ^S^) and the small intrinsically disordered proteins YciG, YmdF and STM14_1829 is compromised in *S.* Typhi. (A) Fold change of *rpoS* and *yciG, STM14_1829* and *ymdF* gene expression at pH 4.5 in STm and STy. *S.* Typhimurium strain that lacks *rpoS, yciG, STM14_1829* and *ymdF* are susceptible to growth inhibition at acidic pH 4.5. (B) Growth of STm wt and STm Δ*rpoS* in buffered LB at pH 4.5. *rpoS* of STy was expressed in STm through allelic exchange, and growth was monitored (C) Growth of STm wt and STm::*rpoS*STy in buffered LB at pH 4.5. (D) Growth of STm wt and STm Δ*rpoS* Δ*yciG* Δ*STM14_1829* Δ*ymdF* in buffered LB at pH 4.5. To generate the growth curves, bacteria were grown in LB-citrate (pH-4.5) or LB-MOPS (pH-7.5) and OD_600_ measured every 15 min at 37°C for 12 h on a Bioscreen C. Typical growth curves representative of three independent experiments are shown. Statistical analysis was performed in GraphPad prism 10.0. Bar graphs represent the mean slopes of exponential growth with error bars representing the standard deviation from three independent assays. *P* values were determined using an un-paired two-tailed *t-*test (B, C & D).

The small intrinsically disordered proteins YciG, YmdF and STM14_1829 have been suggested to play a role in the general stress response and exhibit redundant effects on flagellar synthesis in *S.* Typhimurium (21). Reduced expression of these genes at pH 4.5 in *S.* Typhi is most likely attributable to regulation by σ^38^ (Fig. 4A) (27). *S.* Typhimurium mutants carrying single deletions of *yciG, STM14_1829* or *ymdF* exhibited growth phenotypes similar to that of wildtype *S*. Typhimurium at pH 4.5 (Fig. S2E). Modestly impaired growth of mutant *S.* Typhimurium carrying triple (Δ*yciG* Δ*STM14_1829* Δ*ymdF*) or double gene deletions (Δ*yciG* Δ*ymdF*) was observed, but other double gene deletions (Δ*yciG* Δ*STM14_1829* and Δ*STM14_1829*Δ*ymdF*) did not affect growth at acid pH relative to wildtype *S.* Typhimurium (Fig. S2F). A quadruple *S.* Typhimurium deletion mutant lacking all three intrinsically disordered proteins as well as *rpoS (*Δ*rpoS* Δ*yciG* Δ*STM14_1829*Δ*ymdF)* exhibited reduced growth at pH 4.5 with moderate growth suppression at pH 7.5 compared to wildtype STm (Fig. 4D, Fig. S2G), suggesting that additional σ^38^-dependent loci contribute to acid resistance.

## Discussion

The present study demonstrates reduced survival of *S.* Typhi at acidic pH (pH 4.5) compared to *S.* Typhimurium. This phenotype is biologically relevant, as *Salmonella* is reported to encounter comparable levels of acid stress in the intestinal tract and within the phagolysosomes of phagocytic cells (28–30). We compared the transcriptional response *of S*. Typhi *and S*. Typhimurium at pH 4.5 and identified multiple genetic determinants required for acid stress resistance in *S.* Typhimurium that are deficient in *S.* Typhi.

The acid tolerance response has been extensively studied in *S.* Typhimurium but not in *S.* Typhi (15). Mendoza-Mejía, et al. reported that the deletion of transcriptional regulators (MarR, MarA, MarB), Hya *h*ydrogenase components or fimbrial proteins increase the acid-sensitivity of *S.* Typhi (31). However, they did not investigate the basis for the relative acid sensitivity of *S.* Typhi. In the present study, we measured the genome-wide transcriptional response of *S.* Typhimurium and *S.* Typhi grown in LB broth at neutral (pH 7.5) or acidic (pH 4.5) conditions using RNA-Seq and identified significant differences in the transcriptional response of these two serovars at pH 4.5. While *S.* Typhimurium exhibits the induction of numerous acid-responsive genes, many orthologous genes are either suppressed or pseudogenized in *S.* Typhi.

An extensive literature describes the importance of amino acid decarboxylases in the acid resistance of *Salmonella*, which convert amino acids into corresponding amines to maintain cytosolic pH. The arginine and lysine decarboxylation systems, encoded by *cadA* and *adiC* respectively, have been implicated in the acid response of *Salmonella* (23) (24). Consistent with previous findings, we observed *cadA* and *adiC* induction in both *S.* Typhimurium and *S.* Typhi under acidic conditions. Deletion of these genes decreased growth at acidic pH (data not shown), suggesting a conserved acid resistance mechanism in both serovars. In *S.* Typhimurium, despite the pH-dependent induction of *speF* and *speC*, which encode the ornithine decarboxylase system, deletion of these genes did not affect survival at pH 4.5 (data not shown). In *S.* Typhi, the ornithine decarboxylase system is non-functional due to pseudogenization of the *speC* gene and a lack of induction of *speF* expression (Table S1).

We investigated the role of membrane proteins in the cellular stress response, as they maintain homeostasis by acting as sensors for downstream signaling cascades and transporters of substrates across the bacterial membrane (32). Outer membrane proteins (OMPs) contribute to stress responses in enteric bacteria including *E. coli, Yersinia,* and *Shigella* and may exhibit up- or down-regulation under environmental stress (33). OmpC and OmpF have been reported to be required for acid stress resistance in *E. coli* (34). However, PhoE was the most strongly induced OMP in *S.* Typhimurium at pH 4.5 and is a pseudogene in *S.* Typhi (Fig. 3C). Deletion mutants of *ompC, ompD, ompF* and *phoE* were created in *S.* Typhimurium, and growth curve assays demonstrated the essentiality of *ompC* and *phoE* for *S.* Typhimurium growth at pH 4.5 (Fig. 3A). Deficient *ompC* expression and pseudogenization of *phoE* in *S.* Typhi appear to contribute to the deficient acid resistance of *S.* Typhi. In addition, expression of the membrane-bound periplasmic-facing hydrogenase (*hyd*) operon was observed in *S.* Typhimurium at pH 4.5, but *hydB* is a pseudogene in *S.* Typhi. This aerobic hydrogenase converts protons to molecular hydrogen, which raises cytoplasmic pH (35). Introducing the cloned *S.* Typhimurium *phoE* or *hydB* genes into *S.* Typhi partially restored growth at pH 4.5, confirming the contribution of these pseudogenes to the acid sensitivity of *S.* Typhi.

Comparison of transcriptional regulator expression in *S.* Typhimurium and *S.* Typhi at acid and neutral pH did not reveal differential expression of known key regulators of acid stress resistance, including *phoPQ, fur, envZ* and *ompR* (12, 13, 36). However, *rpoS* (σ^S^, σ^38^) is known to be non-functional in *S.* Typhi Ty2, as well as in approximately one-third of natural *S.* Typhi isolates (37), and has been shown to regulate the acid tolerance response in *S.* Typhimurium (14). RpoS inactivation impaired growth of *S*. Typhimurium at pH 4.5. However, despite the non-functional *rpoS* in the S. Typhi Ty2 strain, no significant difference in growth at pH 4.5 was observed when *S.* Typhi CT18 carrying a functional *rpoS* allele was compared with *rpoS-*deficient *S.* Typhi Ty2 (Fig. S2B).

Lastly, our transcriptional analysis revealed the upregulation of intrinsically disordered protein (IDP)-encoding genes *yciG, STM14_1829* and *ymdF* in *S.* Typhimurium with no induction detected in *S.* Typhi. The three genes exhibit a high degree of sequence similarity (80% identity), and their encoded proteins are rich in conserved, repeated hydrophilic motifs. The IDPs have been previously implicated in stress responses and flagella-dependent motility of *S.* Typhimurium (21, 23). RpoS is known to regulate the expression of these IDPs during acid stress (27). In *S.* Typhimurium, the deletion of *yciG*, *STM14_1829,* or *ymdF* did not increase the growth impairment of a Δ*rpoS* mutant strain. However, the deletion of *rpoS* and all three IDP genes (Δ*rpoS* Δ*yciG* Δ*STM14_1829* Δ*ymdF*) resulted in a greater growth impairment at pH 4.5, suggesting a contribution of additional RpoS-dependent genes to acid resistance and redundancy of function among the IDPs (Fig. 4D).

Collectively, these observations show that the reduced acid stress resistance of *S.* Typhi in comparison to *S.* Typhimurium is attributable to the contribution of multiple genetic differences, including genes encoding outer membrane proteins, a periplasmic hydrogenase, the transcriptional regulator RpoS, and stress response IDPs, which are absent or dysregulated in *S.* Typhi. Most importantly, the loss of acid-resistance in *S.* Typhi by genomic decay raises the question of why this human-specific serovar no longer requires the ability to grow at pH 4.5. Part of the explanation may be the higher pH of the human large intestine in comparison to mice, the natural host of *S.* Typhimurium (38, 39). In addition, the loss of acid resistance suggests that acidification of the *S.* Typhi-containing vacuole in human phagocytes may be impaired compared to that of *S.* Typhimurium, a hypothesis that is currently being investigated.

## MATERIALS AND METHODS

### Bacterial strains and growth conditions

Bacterial strains used in this study are listed in Table S2. Bacteria were routinely grown in LB-Miller broth containing 10 g/L tryptone, 5 g/L yeast extract, and 10 g/L NaCl. LB agar contained 15 g/L Bacto Agar. For experiments, bacteria were grown in 25 mL media in 250 mL Erlenmeyer flasks, unless otherwise indicated. When required, the antibiotic kanamycin (Km) was added to a final concentration of 50 μg/mL, ampicillin (Amp) to 100 μg/mL, and tetracycline (Tet) to 25 μg/mL. For pH experiments, pH of LB was adjusted using HCl or NaOH and buffered with sodium citrate (pH 3.0-4.5), MES (pH 5.0-6.0) or MOPS (pH 6.5-8.0) at a 100 mM final concentration. Unbuffered LB-Miller broth (LB-reg) had a pH of 6.8-6.9.

### RNA isolation, cDNA library preparation and Illumina sequencing

Total RNA from *S.* Typhi Ty2 or *S*. Typhimurium 14028s grown at acidic pH 4.5 or neutral pH 7.5 was isolated using TRIzol and Zymo Kits. Briefly, 1 mL of overnight bacterial culture grown in LB-reg was washed twice in either LB-citrate pH 4.5 or LB-MOPS pH 7.5 then diluted 1:100 into 25 mL of LB at the respective pH in 250 mL sterilized Erlenmeyer flasks. Cells were grown at 37°C with shaking at 200 rpm until the OD_600_ reached 0.3. Six mL of the cultures were mixed with 2/5th volume of ice-cold RNA-stabilizing solution (containing 5% acidic phenol and 95% ethanol) and incubated on ice for 30 min to stabilize the RNA (40). Bacteria were then pelleted and immediately solubilized with 1 mL TRIzol, which was immediately used for RNA isolation or stored at −20°C until RNA isolation. The TRIzol homogenate was processed using the Zymo extraction kit and extracted RNA checked for quantity using the Qubit RNA BR assay and quality using the Agilent 2100 Tapestation and RNA assay. A total of 12 RNA samples (three biological replicates of each strain grown in pH 4.5 and pH 7.5) were prepared for this study.

The RNA samples were sent to SeqCenter (Pittsburgh, USA) for library preparation and Illumina sequencing. Samples were DNAse treated with Invitrogen DNAse (RNAse-free). Library preparation was performed using Illumina’s Stranded Total RNA Prep Ligation with Ribo-Zero Plus kit and 10bp unique dual indices (UDI). Sequencing was done on a NovaSeq X Plus, producing paired end 150bp reads. Demultiplexing, quality control, and adapter trimming was performed with bcl-convert (v4.1.5).

### RNA-seq data analysis

The RNA-seq reads were uploaded to the Galaxy web platform and analyzed using the public server at usegalaxy.org (41).The quality of each RNA-seq library was assessed with FastQC v0.12.1 (http://www.bioinformatics.babraham.ac.uk/projects/fastqc/), and Cutadapt v4.4 (https://journal.embnet.org/index.php/embnetjournal/article/view/200) was used for processing. All reads shorter than 20 nucleotides in length after trimming were discarded from further analysis. The remaining reads of each library were aligned to the corresponding genomes (*S.* Typhi 2: NC_004631.1; *S.* Typhimurium 14028s: NC_016856.1 (chromosome) and NC_016855.1 (plasmid)) using STAR v2.7.10b (42), and alignments were filtered with Samtools v1.16.1 (43) using a MAPQ cutoff of 60. The RNA-seq mapping statistics are detailed in Table S3. Reads were assigned to genomic features using featureCounts v2.0.3 (44, 45). DESeq2 (45) was used to identify differentially expressed genes between the two pH conditions in each strain. Biological significance cut-off was set at log^2^FC ≥ 1 or log_2_FC ≤ −1 and p-value < 0.05.

### Measurement of bacterial growth in broth

Bacterial cultures were grown overnight in LB-reg with shaking (200 rpm) at 37°C, then adjusted to OD_600_ ∼1 and diluted 1:100 into LB at respective pH values with pH buffers. Each bacterial suspension was then used to inoculate the wells of a Bioscreen C honeycomb plate (Growth Curves Ltd., Finland), each containing 300 µL of medium, at a 1:10 dilution (i.e. starting OD_600_ of 0.001). The plate was incubated at 37°C with oscillation at 600 rpm for 12 h, with OD_600_ readings taken at 15-min intervals. Wells containing non-inoculated media were included as negative controls. At least three technical replicates and three biological repeats per strain and pH level were performed.

### Construction of deletion mutants in S. Typhimurium 14028s

1. *S.* Typhimurium deletion mutants were constructed using the λ-Red recombineering method (46, 47). Briefly, oligonucleotides listed in Table S4 were used to amplify the Km (kanamycin) resistance cassette from the pKD4 plasmid. The polymerase chain reaction (PCR) product was electroporated into 14028s cells containing the pKD46 or pSIJ8 plasmid. Transformants were selected on LB agar plates containing 50 µg/mL kanamycin at 37°C (for pKD46) or LB agar plates containing 50 µg/mL kanamycin and 100 µg/mL ampicillin at 30°C (for pSIJ8). The Km resistance cassette was removed by FLP-recombinase either by pCP20 or rhamnose induction.

### Construction of pJK770-*hydB*

The *hydB* gene was heterologously expressed in *S.* Typhi Ty2 using the constitutive expression vector pJK770. First, *hydB* was amplified using S. Typhimurium genomic DNA with pJK770 overlapping primers, followed by ligation with pJK770 vector digested with NcoI. The ligated product was transformed into *S.* Typhi Ty2 competent cells through electroporation, and transformants were selected on LB agar plates containing 100 µg/mL ampicillin at 37°C. *S.* Typhi colonies containing the pJK770-*hydB* plasmid was confirmed by Sanger sequencing. pJK770 vector without insert was transformed to generate *S.* Typhi Ty2 pJK770-Vector.

### Allelic exchange

Replacement of S. Typhi Ty2 *t1450* with S. Typhimurium 14028s *STM14_1848* and S. Typhimurium 14028s *rpoS* with S. Typhi Ty2 *rpoS* was performed using the allelic exchange method as described in (48) with some modifications. The *S.* Typhi Ty2 *t1450* gene was first replaced by a *tetRA* cassette using λ-Red recombineering as described previously. Ty2 Δ*t1450*: tetRA was then used as the recipient strain in the allelic exchange protocol. For *rpoS* allelic exchange, the STy2 *rpoS* gene was cloned into the JKe201 *E. coli* strain using pFOK plasmids with 700bp upstream and downstream regions from STm14028s. STm14028s was then used as a recipient strain for allelic exchange with JKe201-pFOK-rpoS.

### Data Availability

RNA-Seq data have been deposited at Gene Expression Omnibus under accession no. GSE317267.

### Statistical analysis

The bacterial growth rate was calculated by performing linear regression analysis on the natural logarithms of OD_600_ values of the exponential growth phase. The slopes of the growth curves were derived from three independent assays, and *P*-values were determined using an unpaired two-tailed *t-*test with *P*<0.05 considered statistically significant.

---

**Fig S1.**
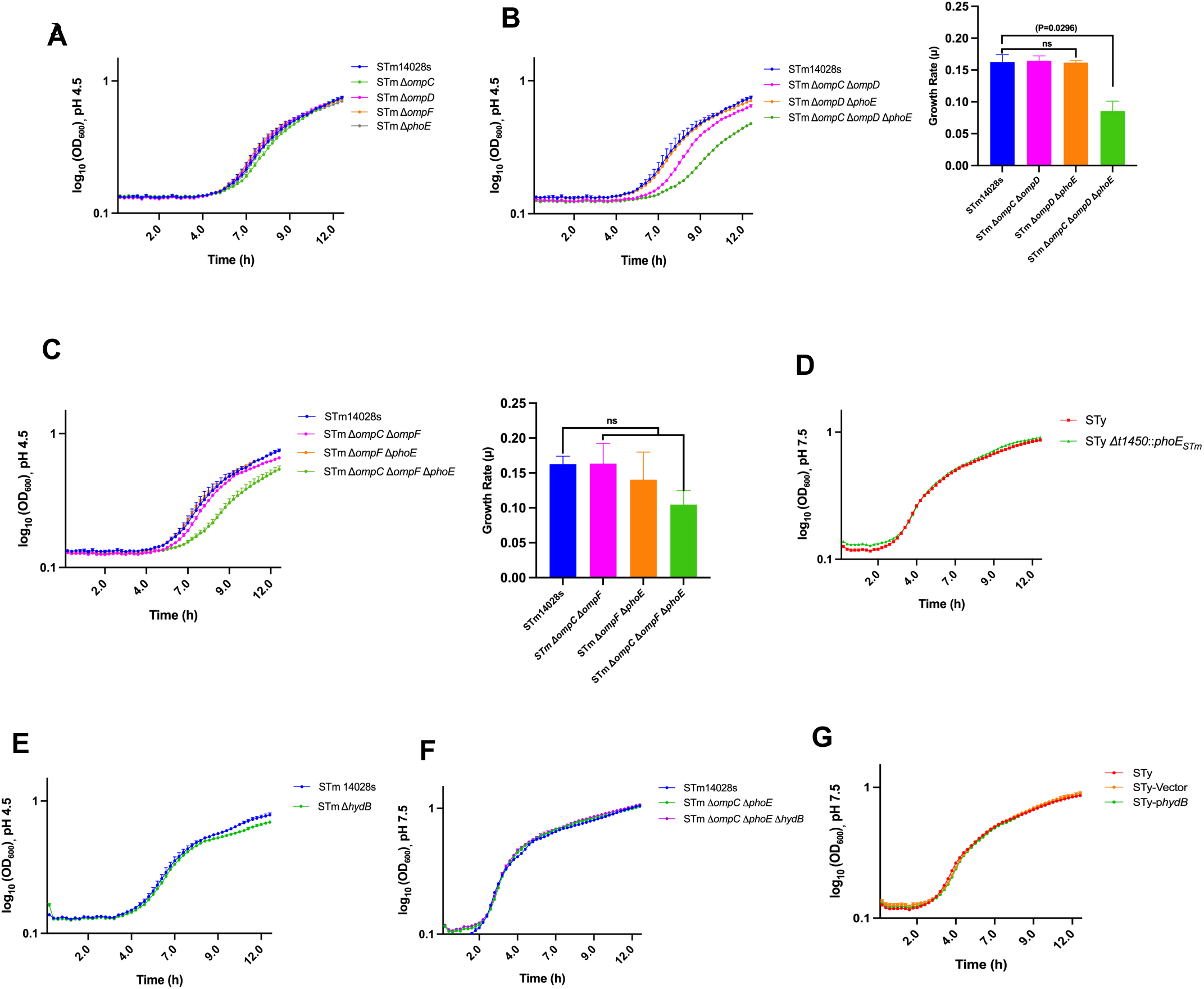
(A) Growth of STm wt, STm Δ*ompC*, STm Δ*ompD*, STm Δ*ompF* and STm Δ*phoE* in buffered LB at pH 4.5. (B) Growth of STm wt, STm Δ*ompC* Δ*ompD*, STm Δ*ompD* Δ*phoE* and STm Δ*ompC* Δ*ompD* Δ*phoE* in buffered LB at pH 4.5. (C) Growth of STm wt, STm Δ*ompC* Δ*ompF*, STm Δ*ompF* Δ*phoE* and STm Δ*ompC* Δ*ompF* Δ*phoE* in buffered LB at pH 4.5 (D) Growth of STm wt, STm Δ*ompC* Δ*phoE* and STm Δ*ompC* Δ*phoE* Δ*hydB* in buffered LB at pH 7.5. (E) Growth of STm wt and STy::*phoE*_STm_ in buffered LB at pH 7.5. (F) Growth of STm wt and STm Δ*hydB* in buffered LB at pH 4.5. (G) Growth of STm wt, STy-Vector (pJK770) and STy-p*hydB* (pJK770-*hydB*) in buffered LB at pH 7.5. No significant growth differences were observed between wildtype and mutant strains of STm and STy at pH 7.5. To generate the growth curves, bacteria were grown in LB-citrate (pH-4.5) and LB-MOPS (pH-7.5) and OD_600_ measured every 15 min at 37°C for 12 h on a Bioscreen C. Statistical analysis was performed in GraphPad prism 10.0. Bar graphs represent the mean slopes of exponential growth with error bars representing the standard deviation from three independent assays. *P* values were determined using an un-paired two-tailed *t-*test (B & C); ns, not significant.

**Fig S2.**
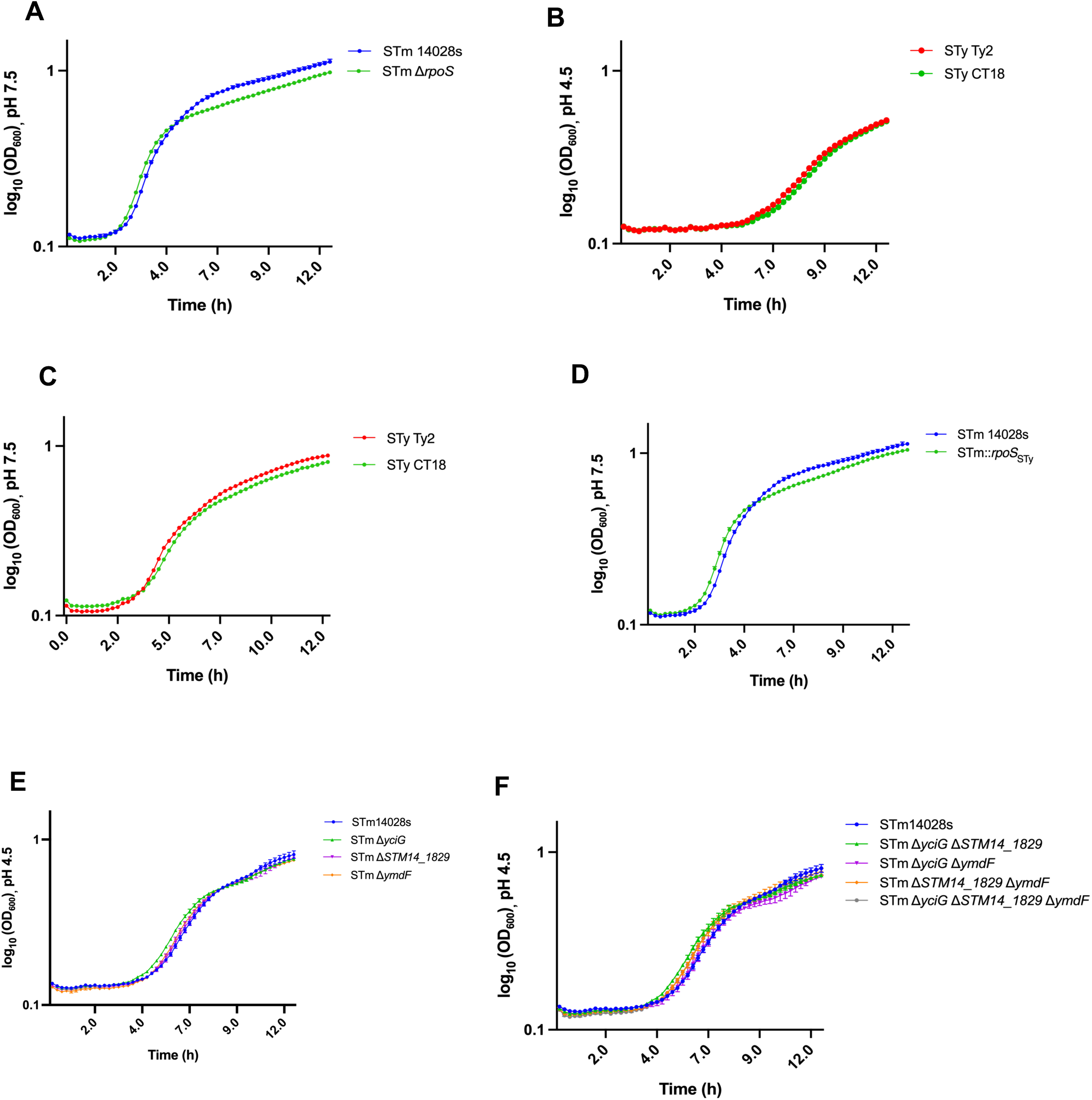
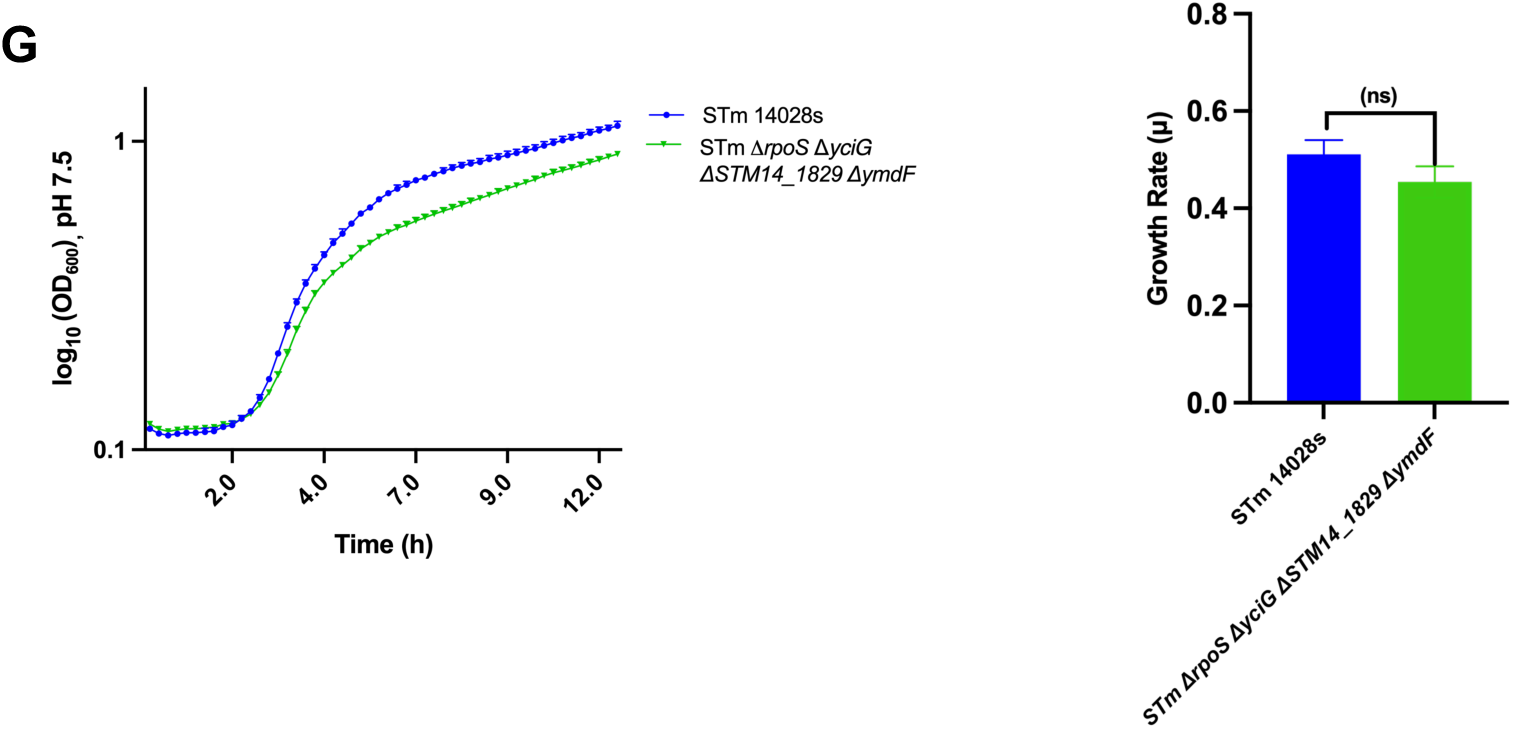
(A) Growth of STm wt and STm Δ*rpoS* in buffered LB at pH 7.5. (B&C) Growth of STy Ty2 and STy CT18 in buffered LB at pH 4.5 & 7.5 respectively. (D) Growth of STm wt and STm::*rpoS*_STy_ in buffered LB at pH 7.5 (E) Growth of STm wt, STm Δ*yciG*, STm Δ*STM14_1829* and STm Δ*ymdF* in buffered LB at pH 4.5. (F) Growth of STm wt, STm Δ*yciG* Δ*STM14_1829*, STm Δ*yciG* Δ*ymdF*, STm Δ*STM14_1829* Δ*ymdF* and STm Δ*yciG* Δ*STM14_1829* Δ*ymdF* in buffered LB at pH 4.5 (G) Growth of STm wt and STm Δ*rpoS* Δ*yciG* Δ*STM14_1829* Δ*ymdF* in buffered LB at pH 7.5. No significant growth differences were observed between wildtype and mutant strains of STm and STy at pH 7.5 except for the quadruple mutant STm Δ*rpoS* Δ*yciG* Δ*STM14_1829* Δ*ymdF,* which showed moderately impaired growth at neutral pH. To generate the growth curves, bacteria were grown in LB-citrate (pH-4.5) or LB-MOPS (pH-7.5) and OD_600_ measured every 15 min at 37°C for 12 h on a Bioscreen C. Typical growth curves representative of three independent experiments are shown. Statistical analysis was performed in GraphPad prism 10.0. Bar graphs represent the mean slopes of exponential growth with error bars representing the standard deviation from three independent assays. *P* values were determined using an un-paired two-tailed *t-*test (G); ns, not significant.

## References

1. Balasubramanian RA, Im J, Lee JA, Jeon HA, Mogeni OA, Kim JA, Rakotozandrindrainy R, Baker SA, Marks FA. The global burden and epidemiology of invasive non-typhoidal *Salmonella* infections. Hum Vaccine Immunother 15:1421–6. 10.1080/21645515.2018.1504717

2. Bhandari J, Thada PK, Hashmi MF, DeVos E. 2025. Typhoid fever. StatPearls. https://www.ncbi.nlm.nih.gov/books/NBK557513/

3. Centers for Disease Control and Prevention. 2013. Incidence and trends of infection with pathogens transmitted commonly through food - foodborne diseases active surveillance network, 10 U.S. sites, 1996-2012. MMWR Morb Mortal Wkly Rep 62:283–7. https://pubmed.ncbi.nlm.nih.gov/23594684/

4. Murray CJ. 1991. Salmonellae in the environment. Rev Sci Tech 10:765–85. 10.20506/rst.10.3.568

5. GBD 2017 Non-Typhoidal Salmonella invasive Disease Collaborators. 2019. The global burden of non-typhoidal *Salmonella* invasive disease: a systematic analysis for the Global Burden of Disease Study 2017. Lancet Infect Dis 19:1312–1324. 10.1016/s1473-3099(19)30418-9

6. Fang FC, Frawley ER, Tapscott T, Vázquez-Torres A. 2016. Bacterial stress responses during host infection. Cell Host Microbe 20:133–43. 10.1016/j.chom.2016.07.009

7. Shen S, Fang FC. 2012. Integrated stress responses in *Salmonella*. Int J Food Microbiol 152:75–81. 10.1016/j.ijfoodmicro.2011.04.017

8. Foster JW, Spector MP. 1995. How *Salmonella* survive against the odds. Annu Rev Microbiol 49:145–74. 10.1146/annurev.mi.49.100195.001045

9. Richard HT, Foster JW. 2003. Acid resistance in *Escherichia coli*. Adv Appl Microbiol 52:167–86. 10.1016/s0065-2164(03)01007-4

10. Alvarez-Ordóñez A, Fernández A, Bernardo A, López M. 2010. Arginine and lysine decarboxylases and the acid tolerance response of *Salmonella* Typhimurium. Int J Food Microbiol 136:278–82. 10.1016/j.ijfoodmicro.2009.09.024

11. Foster JW. 2004. *Escherichia coli* acid resistance: tales of an amateur acidophile. Nat Rev Microbiol 2:898–907. 10.1038/nrmicro1021

12. Bearson BL, Wilson L, Foster JW. 1998. A low pH-inducible, PhoPQ-dependent acid tolerance response protects *Salmonella typhimurium* against inorganic acid stress. J Bacteriol 180:2409–17. 10.1128/jb.180.9.2409-2417.1998

13. Foster JW, Hall HK. 1992. Effect of *Salmonella typhimurium* ferric uptake regulator (*fur*) mutations on iron- and pH-regulated protein synthesis. J Bacteriol 174:4317–23. 10.1128/jb.174.13.4317-4323.1992

14. Lee IS, Lin J, Hall HK, Bearson B, Foster JW. 1995. The stationary-phase sigma factor sigma S (RpoS) is required for a sustained acid tolerance response in virulent *Salmonella typhimurium*. 10.1111/j.1365-2958.1995.mmi_17010155.x

15. Lee IS, Slonczewski JL, Foster JW. 1994. A low-pH-inducible, stationary-phase acid tolerance response in *Salmonella typhimurium*. J Bacteriol 176:1422–6. 10.1128/jb.176.5.1422-1426.1994

16. Tiwari RP, Sachdeva N, Hoondal GS, Grewal JS. 2004. Adaptive acid tolerance *response in Salmonella enterica* serovar *Typhimurium and Salmonella enterica* serovar Typhi. J Basic Microbiol 44:137–46. 10.1002/jobm.200310333

17. Bäumler A, Fang FC. 2013. Host specificity of bacterial pathogens. Cold Spring Harb Perspect Med 3:a010041. 10.1101/cshperspect.a010041

18. Shane AL, Mody RK, Crump JA, Tarr PI, Steiner TS, Kotloff K, Langley JM, Wanke C, Warren CA, Cheng AC, Cantey J, Pickering LK. 2017. 2017 Infectious Diseases Society of America clinical practice guidelines for the diagnosis and management of infectious diarrhea. Clin Infect Dis 65:e45–e80. 10.1093/cid/cix669

19. Sabbagh SC, Forest CG, Lepage C, Leclerc JM, Daigle F.2010. So similar, yet so different: uncovering distinctive features in the genomes of *Salmonella enterica* serovars Typhimurium and Typhi. FEMS Microbiol Lett 305:1–13. 10.1111/j.1574-6968.2010.01904.x

20. Garai P, Gnanadhas DP, Chakravortty D. 2012. *Salmonella enterica* serovars Typhimurium and Typhi as model organisms: revealing paradigm of host-pathogen interactions. Virulence 3:377–88. 10.4161/viru.21087

21. Oguri T, Kwon Y, Woo JKK, Prehna G, Lee H, Ning M, Won KJ, Lee J, Mei S, Shi Y, Jeong H, Lee H. 2018. A family of small intrinsically disordered proteins involved in flagellum-dependent motility in *Salmonella enterica*. J Bacteriol 201:e00415–18. 10.1128/jb.00415-18

22. Robbe-Saule V, Norel F. 1999. The rpoS mutant allele of *Salmonella* typhi Ty2 is identical to that of the live typhoid vaccine Ty21a. FEMS Microbiol Lett 170:141–3. 10.1111/j.1574-6968.1999.tb13366.x

23. Ryan D, Pati NB, Ojha UK, Padhi C, Ray S, Jaiswal S, Singh GP, Mannala GK, Schultze T, Chakraborty T, Suar M. 2015. Global transcriptome and mutagenic analyses of the acid tolerance response of *Salmonella enterica* serovar Typhimurium. Appl Environ Microbiol 81:8054–65. 10.1128/aem.02172-15

24. Viala JP, Méresse S, Pocachard B, Guilhon AA, Aussel L, Barras F. 2011. Sensing and adaptation to low pH mediated by inducible amino acid decarboxylases in *Salmonella*. PLoS One 6:e22397. 10.1371/journal.pone.0022397

25. Boyd EF, Porwollik S, Blackmer F, McClelland M. 2003. Differences in gene content among *Salmonella enterica* serovar Typhi isolates. J Clin Microbiol 41:3823–8. 10.1128/jcm.41.8.3823-3828.2003

26. Parkin A, Bowman L, Roessler MM, Davies RA, Palmer T, Armstrong FA, Sargent F. 2012. How *Salmonella* oxidises H(2) under aerobic conditions. FEBS Lett 586:536–44. 10.1016/j.febslet.2011.07.044

27. Lago M, Monteil V, Douche T, Guglielmini J, Criscuolo A, Maufrais C, Matondo M, Norel F. 2017. Proteome remodelling by the stress sigma factor RpoS/σ(S) in *Salmonella*: identification of small proteins and evidence for post-transcriptional regulation. Sci Rep 7:2127. 10.1038/s41598-017-02362-3

28. Rathman M, Sjaastad MD, Falkow S. 1996. Acidification of phagosomes containing *Salmonella typhimurium* in murine macrophages. Infect Immun 64:2765–73. 10.1128/iai.64.7.2765-2773.1996

29. McConnell EL, Basit AW, Murdan S. 2008. Measurements of rat and mouse gastrointestinal pH, fluid and lymphoid tissue, and implications for in-vivo experiments. J Pharm Pharmacol 60:63–70 10.1211/jpp.60.1.0008

30. Ibarra JA, Steele-Mortimer O. 2009. *Salmonella*--the ultimate insider. *Salmonella* virulence factors that modulate intracellular survival. Cell Microbiol 11:1579–86. 10.1111/j.1462-5822.2009.01368.x

31. Mendoza-Mejía BD, Medina-Aparicio L, Serrano-Fujarte I, Vázquez A, Calva E, Hernández-Lucas I.2021. *Salmonella enterica* serovar Typhi genomic regions involved in low pH resistance and in invasion and replication in human macrophages. Ann Microbiol 71. 10.1186/s13213-021-01629-5

32. Walsh NP, Alba BM, Bose B, Gross CA, Sauer RT. OMP peptide signals initiate the envelope-stress response by activating DegS protease via relief of inhibition mediated by its PDZ domain. Cell 113:61–71. 10.1016/s0092-8674(03)00203-4

33. Harikrishnan H, Ismail A, Banga Singh KK. 2013. Temperature-regulated expression of outer membrane proteins in *Shigella flexneri*. Gut Pathog 5:38. 10.1186/1757-4749-5-38

34. Bekhit A, Fukamachi T, Saito H, Kobayashi H. 2011. The role of OmpC and OmpF in acidic resistance in *Escherichia coli*. Biol Pharm Bull 34:330–4. 10.1248/bpb.34.330

35. Zbell AL, Benoit SL, Maier RJ. 2007. Differential expression of NiFe uptake-type hydrogenase genes in *Salmonella enterica* serovar Typhimurium. Microbiology (Reading) 153:3508–3516. 10.1099/mic.0.2007/009027-0

36. Bang IS, Kim BH, Foster JW, Park YK. 2000. OmpR regulates the stationary-phase acid tolerance response of *Salmonella enterica* serovar typhimurium. J Bacteriol 182:2245–52. 10.1128/jb.182.8.2245-2252.2000

37. Robbe-Saule V, Algorta G, Rouilhac I, Norel F. 2003. Characterization of the RpoS status of clinical isolates of *Salmonella enterica*. Appl Environ Microbiol 69:4352–8. 10.1128/aem.69.8.4352-4358.2003

38. Yamamura R, Inoue KY, Nishino K, Yamasaki S. 2023. Intestinal and fecal pH in human health. Front Microbiomes 2:1192316. https://www.frontiersin.org/journals/microbiomes/articles/10.3389/frmbi.2023.1192316

39. Fallingborg J. 1999. Intraluminal pH of the human gastrointestinal tract. Dan Med Bull 46:183–96. https://pubmed.ncbi.nlm.nih.gov/10421978/

40. Tedin K, Bläsi U. 1996. The RNA chain elongation rate of the lambda late mRNA is unaffected by high levels of ppGpp in the absence of amino acid starvation. J Biol Chem 271:17675–86. 10.1074/jbc.271.30.17675

41. Anonymous. 2024. The Galaxy platform for accessible, reproducible, and collaborative data analyses: 2024 update. Nucleic Acids Res 52:W83–W94. 10.1093/nar/gkae410

42. Dobin A, Davis CA, Schlesinger F, Drenkow J, Zaleski C, Jha S, Batut P, Chaisson M, Gingeras TR. 2013. STAR: ultrafast universal RNA-seq aligner. Bioinformatics 29:15–21. 10.1093/bioinformatics/bts635

43. Danecek P, Bonfield JK, Liddle J, Marshall J, Ohan V, Pollard MO, Whitwham A, Keane T, McCarthy SA, Davies RM, Li H. 2021. Twelve years of SAMtools and BCFtools. Gigascience 10:giab008. 10.1093/gigascience/giab008

44. Liao Y, Smyth GK, Shi W. 2014. featureCounts: an efficient general purpose program for assigning sequence reads to genomic features. Bioinformatics 30:923–30. 10.1093/bioinformatics/btt656

45. Love MI, Huber W, Anders S. 2014. Moderated estimation of fold change and dispersion for RNA-seq data with DESeq2. Genome Biol 15:550. 10.1186/s13059-014-0550-8

46. Datsenko KA, Wanner BL. 2000. One-step inactivation of chromosomal genes in *Escherichia coli* K-12 using PCR products. Proc Natl Acad Sci U S A 97:6640–5. 10.1073/pnas.120163297

47. Jensen SI, Lennen RM, Herrgård MJ, Nielsen AT. 2015. Seven gene deletions in seven days: Fast generation of *Escherichia coli* strains tolerant to acetate and osmotic stress. Sci Rep 5:17874. 10.1038/srep17874

48. Cianfanelli FR, Cunrath O, Bumann D. 2020. Efficient dual-negative selection for bacterial genome editing. BMC Microbiol 20:129. 10.1186/s12866-020-01819-2

